# EEG Responses to Auditory Figure-Ground Perception

**DOI:** 10.1101/2022.03.03.482346

**Authors:** Xiaoxuan Guo, Pradeep Dheerendra, Ester Benzaquén, William Sedley, Timothy D Griffiths

## Abstract

Speech-in-noise difficulty is commonly reported among hearing-impaired individuals. Recent work has established generic behavioural measures of sound segregation and grouping that are related to speech-in-noise processing but do not require language. In this study, we assessed potential clinical electroencephalographic (EEG) measures of central auditory grouping (stochastic figure-ground test) and speech-in-noise perception (speech-in-babble test) with and without relevant tasks. Auditory targets were presented within background noise (16 talker-babble or randomly generated pure-tones) in 50% of the trials and composed either a figure (pure-tone frequency chords repeating over time) or speech (English names). EEG was recorded while participants were presented with the target stimuli (figure or speech) under different attentional states (relevant task or visual-distractor task). EEG time-domain analysis demonstrated enhanced negative responses during detection of both types of auditory targets within the time window 650-850 ms but only figure detection produced significantly enhanced responses under the distracted condition. Further single-channel analysis showed that simple vertex-to-mastoid acquisition defines a very similar response to more complex arrays based on multiple channels. Evoked-potentials to the generic figure-ground task therefore represent a potential clinical measure of grouping relevant to real-world listening that can be assessed irrespective of language knowledge and expertise even without a relevant task.

## Introduction ^1^

Speech perception is often challenged with competing speech sounds (e.g., multiple speakers talking simultaneously) or environmental sounds (e.g., air conditioning system, traffic noise, etc.). Difficulty understanding speech is often described as “speech-in-noise” difficulty, or more colloquially, the “cocktail party problem” (Cherry, 1953). Speech-in-noise (SiN) perception is not only essential for people to perform their daily social and occupational commitments; as with other types of hearing impairment, having difficulty understanding speech could also lead to isolation, psychiatric disorders such as depression and anxiety disorder, and overall lower quality of life (Rutherford et al., 2018; Scinicariello et al., 2019; Blazer & Tucci, 2019); the underlying mechanisms for SiN is also considered a potential factor that accounts for the association between hearing loss and development of later-life dementia (Griffiths et al., 2020).

SiN tests are considered a powerful behavioural measure for real-world listening difficulties. Unlike pure tone audiometry (PTA) test, SiN tests capture not only defective peripheral hearing but also central auditory grouping, auditory working memory, language competence, and other predictors of auditory cognition (Lad et al., 2020; Holmes & Griffiths, 2019; Skoe & Karayanidi, 2019). While SiN stimuli are considered ecological, their linguistic content also means that optimal effects can only be obtained from a specific group of people (e.g., educated English native speakers with a particular accent), without wider generalisability. The linguistic or social cues embedded in the stimuli could also help patients generate expectations and thus compensate for compromised auditory processing mechanisms. To address this limitation, Stochastic Figure-Ground (SFG), a prototype for SiN testing that can be more widely applicable (e.g. children or speakers of any language), has been developed (Holmes et al., 2021; Holmes & Griffiths, 2019; Teki et al., 2011). SFG consists of a set of tones of multiple frequencies repeating over time (figure) against a background of pure-tone segments randomised over frequency and time (ground) (Teki et al., 2011). Extraction of the “figure” requires successful segregation based on perceptual commonalities as well as a sequential grouping in time-frequency space, which is similar to tracking speech targets with background noise. Previous work has shown that participants can successfully detect figures, and that performance improves with increasing figure *coherence* (Teki et al., 2013, 2016; Holmes & Griffiths, 2019), which refers to the number of spectral elements that repeat over time. Neural imaging studies also discovered that SFG engages high-level mechanisms, some of which are not within traditional auditory areas, including the superior temporal sulcus (STS) bilaterally, the intraparietal sulcus (IPS) and the planum temporale (PT), indicating that auditory grouping does not only involve processes in the early auditory cortices (Teki et al., 2011). Source analysis with electroencephalography (EEG) also found that object-related negativity (ORN) elicited by SFG were generated in the superior temporal gyrus (STG), IPS, the cingulate gyrus, as well as some frontal regions (Alain et al., 2001; Arnott et al., 2011; Tóth et al., 2016).

While previous psychophysical and neuroimaging studies have detailed behavioural and neural responses to SFG, clinical applications of these protocols have not been developed. The administration of elaborate testing protocols or expensive neuroimaging techniques is impractical for clinical settings. To develop a hearing test for central auditory grouping with simple active tasks and robust and accessible brain recordings in audiology clinics, we assessed the effectiveness of using a single EEG electrode montage similar to that used for brain-stem auditory evoked potential (BSAEPs) while carrying out psychophysical tasks. The data demonstrate a vertex response with a delay of greater than 100 ms that can be recorded both in the presence and absence of a relevant task. The results suggest that SFG could provide useful clinical measures of real-world listening ability in patients without having to perform a behavioural task. We also examined ERP responses to a SiN test, from the vertex, which were similar to the SFG evoked responses, but less robust, and not present without an active auditory task.

## 1. Materials and Methods

### 1.1 Participants

A total of 18 participants (4 male) aged 18 to 53 (mean ± SD: 25.47±10.57) of both sexes were recruited for the study. Audiometric thresholds were measured and recorded in decibels hearing level (dB HL) for each participant before the main experiment (Figure 1) and only people with clinically normal hearing thresholds were included in the study (seven frequencies averaged lower than 20dB HL in either ear). Participants had no history of auditory disorders (e.g., auditory processing disorders, misophonia, or tinnitus), neurological disorders or traumatic brain injuries, and were not taking psychotropic drugs or medication. Experimental procedures were approved by the research ethics committee of Newcastle University and written informed consent was obtained from all participants.

**Figure 1.**
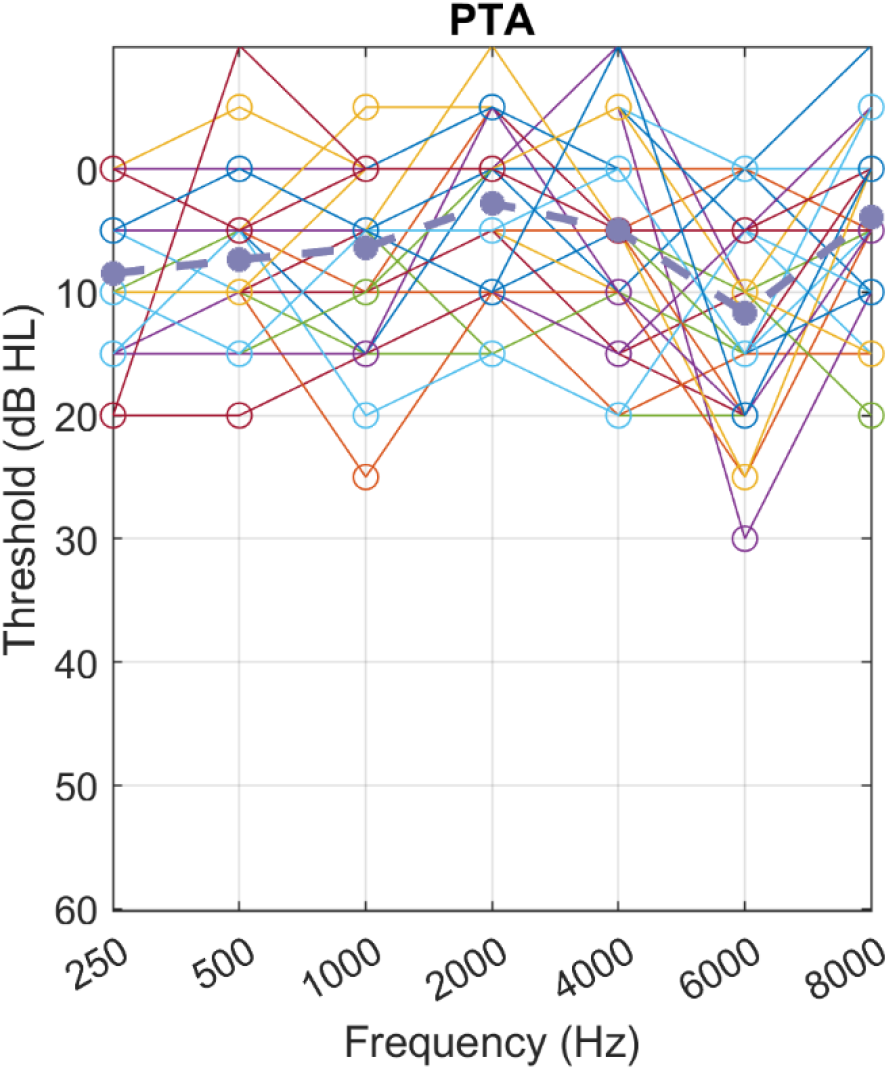
Pure-tone audiograms for the participants. The dashed line shows the group mean.

### 1.2 Stimuli

The auditory stimuli were based on the SiN test used by Holmes & Griffiths (2019) and the SFG stimuli developed by Teki et al. (2011). Each stimulus comprised a sequence of random chords with 15 pure tone components per chord and a 50 ms duration with 0 ms inter-chord interval. Each stimulus contained two segments; the first segment lasted for 500 ms and was ground-only, while the second segment, also 500 ms long, was divided into two conditions: condition one presented a figure (coherence=6, 50% of the trials), condition two contained no figure (coherence=0, 50% of the trials). Coherence of 6 has been shown to elicit high detection sensitivity previously so the figure used here is considered highly coherent (Teki et al., 2013). The speech-in-noise stimuli consisted of English names spoken in a British accent and 16-talker babble noise. Similar to the SFG stimulus design, SiN also contained two segments, with the first being only babble noise lasting for about 500 ms and the second with either 50% trials of babble noise or 50% trials of speech (SNR= −3 dB) amidst babble noise. Auditory stimulus onset for both SFG and SiN is defined as 0 ms, and figure onset as 500 ms. A distractor visual task was adopted from the Random Dot Kinematograms (RDK) test (Fleming et al., 2018), where white dots were presented on grey background with a fixation spot at the centre of the screen. The size of the dots was 0.12 degrees (deg) diameter, and they moved at a speed of 5 deg/sec with a density of 30 dots/deg^2^. The first segment of RDK was 500ms of random movement. Again, the second segment was divided into two conditions: the first condition had motion coherence of 0.5, creating coherent motion to either the left or right. The coherent condition accounts for 80% of the trials, and the rest of the trials belonged to the random-movement condition, which had motion coherence of 0.

### 1.3 Procedure

The experiment was carried out in a sound-proof booth. Stimuli were presented using headphones (Sennheiser HD 380 Pro) connected to an external sound card (RME FireFace UC). Participants were asked to sit in front of the LCD display (Dell Inc.) in the booth with their eyes about 1 metre away from the screen.

The experiment contained two blocks, first the distractor block and then the active block to reduce participants’ learning of the generic properties and structure of the stimuli before doing the active task. During the distractor task, participants were instructed to fixate on the screen and press a key if there is no coherent motion of dots in the RDK task while ignoring the SFG or SiN stimuli during the distractor block. Participants were also shown the visual distractors in the active block, but they were asked to ignore the moving dots and fixate on the fixation point at the centre of the screen and respond when there was no figure or no speech present for the SFG or SiN tasks. The SFG and SiN trials were randomly interleaved, and the inter-trial interval was 1.3 s (1.1-1.5s). The trail length was 2.3s in total, and there were 200 SFG trials and 200 SiN trials in each block, making 800 trials in total.

### 1.4 Data Acquisition and Analysis

The behavioural response was analysed with a measure of detection sensitivity: d prime (d’). The d’ was calculated as the difference of standardised hit rate and false alarm rate (d’ = z(H) - z(F)). The extreme values were adjusted by replacing 0 with 0.5/trial number, and 1 with (trial number−0.5)/trial number (Macmillan & Kaplan, 1985). Separate d’ were calculated for SFG and SiN stimuli and for active and distractor tasks. Correlation was performed to check the relationship between PTA and the behavioural as well as neurophysiological measures.

EEG data were acquired using a 128-channel BioSemi system. MATLAB R2021a with EEGLAB version 2019 was used to preprocess the EEG data. Data analysis was carried out with multiple channels as well as with just one channel that can be carried out in clinics (the vertex, A1). For the multiple-channel analysis, the original sampling rate of 2048 Hz was reduced by a factor of 8 to 256 Hz in order to increase the processing speed. The continuous EEG data were filtered from 0.1—30 Hz using a highpass Infinite Impulse Response Butterworth filter and then a lowpass band-pass Butterworth filter. The Artifact subspace reconstruction (ASR) tool was used to detect noisy channels: channels poorly correlated (r<0.6) with their random sample consensus reconstruction were rejected and interpolated (8.58 ± 3.67). If over 10% of channels were rejected, the participant was removed from further analysis. This resulted in the rejection of one participant. The data were re-referenced to the common average and epoched from −200 to 1000 ms with a baseline set at 400-500 ms, which is 100 ms before the stimulus onset. Independent component analysis (ICA) was conducted, and components constituting eye artefacts were rejected via visual inspection. Trial rejection was performed based on probability (>5 SD) and kurtosis (>8). To reduce data loss due to the high montage during trial rejection, temporarily noisy channels were identified and interpolated on a trial-by-trial basis before trial rejection: if a channel exceeded a voltage of 100 mV in a given trial, this channel would be interpolated on that trial only; if more than 3 channels were identified on a given trial, this trial would be rejected from analysis. Event-related potentials (ERPs) were computed across all good trials and across the vertex and selected neighbouring electrodes (A1, B1, C1, D1, D15, A2). To calculate the difference at sensor level in the time domain between two conditions, Monte Carlo permutation testing was used at the 500-1000ms time window (corresponding to the figure/speech stimulus) with 1000 iterations and at 0.025 false alarm rate. Cluster correction (threshold at *p* < 0.05) was also performed to avoid multiple comparisons problem across time points and channels. Scalp maps were plotted with cluster-based permutation test across all electrodes at two time windows (600 - 800 ms and 800 - 1000 ms).

For clinical use, after down-sampling and filtering, three channels (A1, D32, B10) were selected for the single-channel analysis. D32 and B10 were used to re-reference the data as substitutes for the mastoids. They are located at a similar position as P9 and P10 in a 64-channel system just behind the ears. Similar to the multi-channel analysis, probability of 5 and kurtosis of 8 were used to clean up trials with artefacts. The preprocessed data were then epoched from –X to Y, timelocked to sound onset and ERPs were computed across all good trials at the vertex (channel A1, equivalent to Cz). The amplitude at the vertex over both defined time windows (600 - 800 and 800 - 1000) was averaged during the active and distractor tasks for the SFG and SiN conditions separately. The amplitude difference between figure and ground, and speech and noise were calculated per participant. A two-way repeated measures Analysis of Variance (ANOVA) was also performed to examine the two within-subject factors, ‘Stimulus Type’ (SiN vs. SFG) and ‘Condition’ (active vs. distractor) and their interaction.

## 2. Results

The behavioural results show an average d’ of around 2~3 for all four tasks (see Table 1). Based on the mean statistics, the SFG task elicited a similar detection sensitivity to the SiN task (t (11) = 0.733, p=0.473, Cohen’s d=0.168). Pure-tone audiograms did not correlate with d’ or the EEG amplitudes (ps>0.50).

**Table 1.** Detection sensitivity (d’) for SFG, SiN and distractor visual tasks. Final row shows the means and standard deviations.

| Participant | Active SFG | Active SiN | Distractor visual |
| --- | --- | --- | --- |
| 1 | 3.609 | 3.981 | 3.941 |
| 2 | 3.804 | 5.152 | 5.083 |
| 3 | 2.275 | 2.362 | 3.196 |
| 4 | 2.926 | 2.592 | 4.187 |
| 5 | 2.048 | 2.760 | 4.044 |
| 6 | 3.656 | 3.156 | 2.804 |
| 7 | 3.335 | 0.869 | 3.858 |
| 8 | 3.156 | 2.247 | 3.400 |
| 9 | 2.485 | 3.751 | 1.456 |
| 10 | 2.412 | 2.109 | 3.334 |
| 11 | 2.926 | 3.981 | 2.849 |
| 12 | 3.981 | 3.459 | 4.084 |
| 13 | 4.107 | 3.609 | 3.553 |
| 14 | 3.417 | 3.804 | 3.497 |
| 15 | 1.555 | 0.892 | 1.857 |
| 16 | 2.926 | 2.745 | 1.916 |
| 17 | 3.156 | 1.831 | 1.858 |
| 18 | 2.327 | 2.276 | 2.234 |
| Mean (SD) | 3.004 (0.690) | 2.849 (1.082) | 3.131 (0.984) |

### 2.1 Multi-channel ERP Topographic Analysis

When inspecting across all channels, central channels showed significantly stronger responses. The scalp maps of figure and ground, speech and noise, and the differences at 600-800 ms and 800-1000 ms averaged over time are shown in Figure 2. For SFG, the negativity was mostly driven by fronto-central channels, whereas for SiN, the distribution is relatively widespread, and more posterior compared to SFG. A similar topographic distribution of SFG was observed for both conditions at both time windows, but the distractor condition only showed significant differences between figure and ground at the later time window. The SiN task, however, showed no significant differences between the speech and noise stimuli across channels.

**Figure 2.**
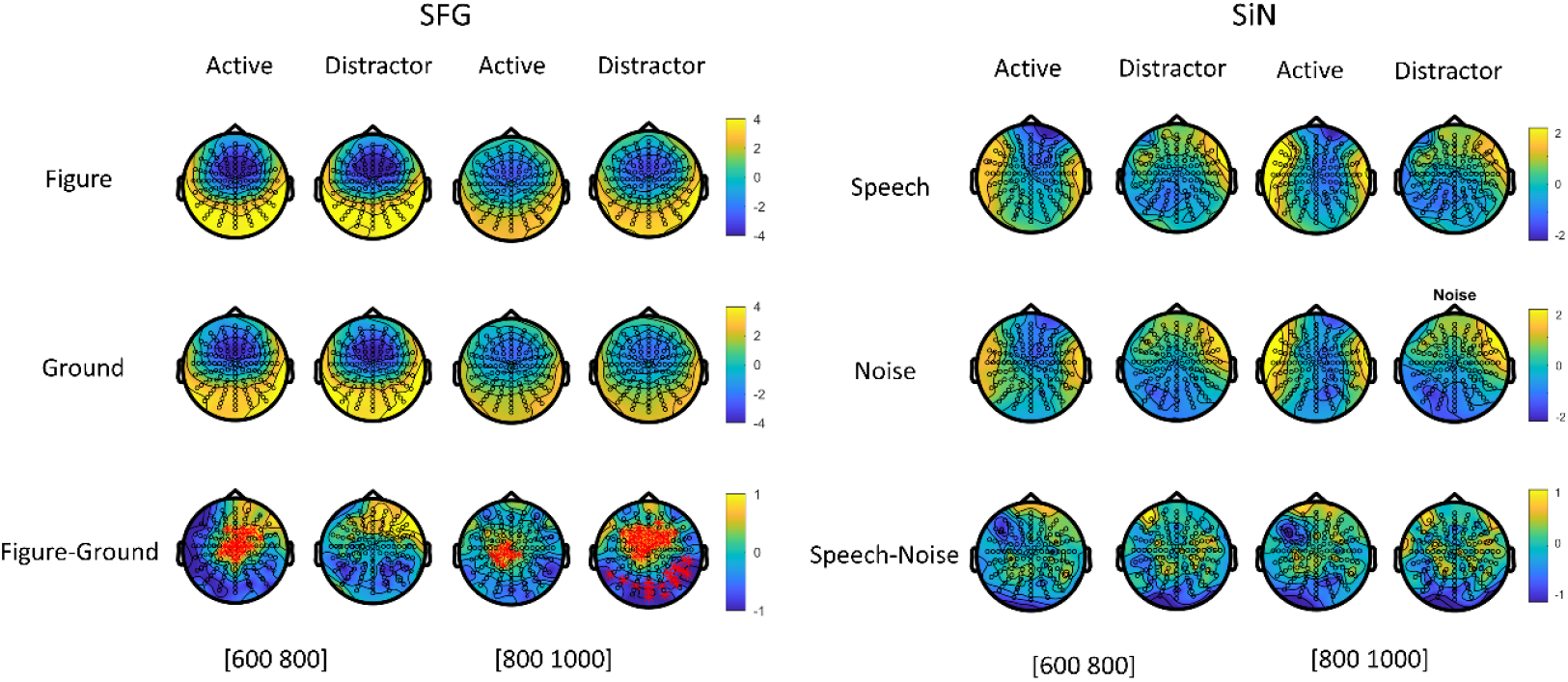
Topographic maps of SFG and SiN of the active and distractor condition at 600 - 800 ms and 800 −1000 ms. The bottom panel shows amplitude differences between figure and ground, and speech and noise (calculated as figure minus ground and speech minus noise). Channels that generated significant voltage differences are highlighted in red (p < 0.05, *cluster-corrected*).

### 2.2 Single-Channel Time-Locked Analysis

The ERP grand averages for the active and distractor SFG and SiN are illustrated in (Figure 3). Through visual inspection, all task conditions showed robust N1 responses to the auditory stimuli. A clear separation elicited by the auditory target from the background was demonstrated post-stimulus onset (i.e., 500 ms) for both SFG and SiN tasks. The auditory targets (figure and speech) elicited greater negativity than the background (ground and noise) alone. Figure tracking started to show significantly enhanced negativity compared to the ground upon the onset of the auditory targets in both active and distractor conditions (approximately 139 ms), peaked around 300 ms after figure onset, and reached statistical significance (p<0.05, cluster-corrected) for about 266 ms for both conditions. Such effect was only significant in the figure-ground paradigm, whilst the speech-in-noise paradigm merely elicited a comparable trend. Speech did display significantly greater amplitude in the active condition at 445 ms post-stimulus onset and lasted for 55 ms (p<0.05, cluster-corrected), in the active condition only. This was in the opposite direction to other differences seen, and we interpret this as a rebound overshoot following the initial figure or speech-related negative potential. A similar trend was seen in the active SFG condition.

**Figure 3.**
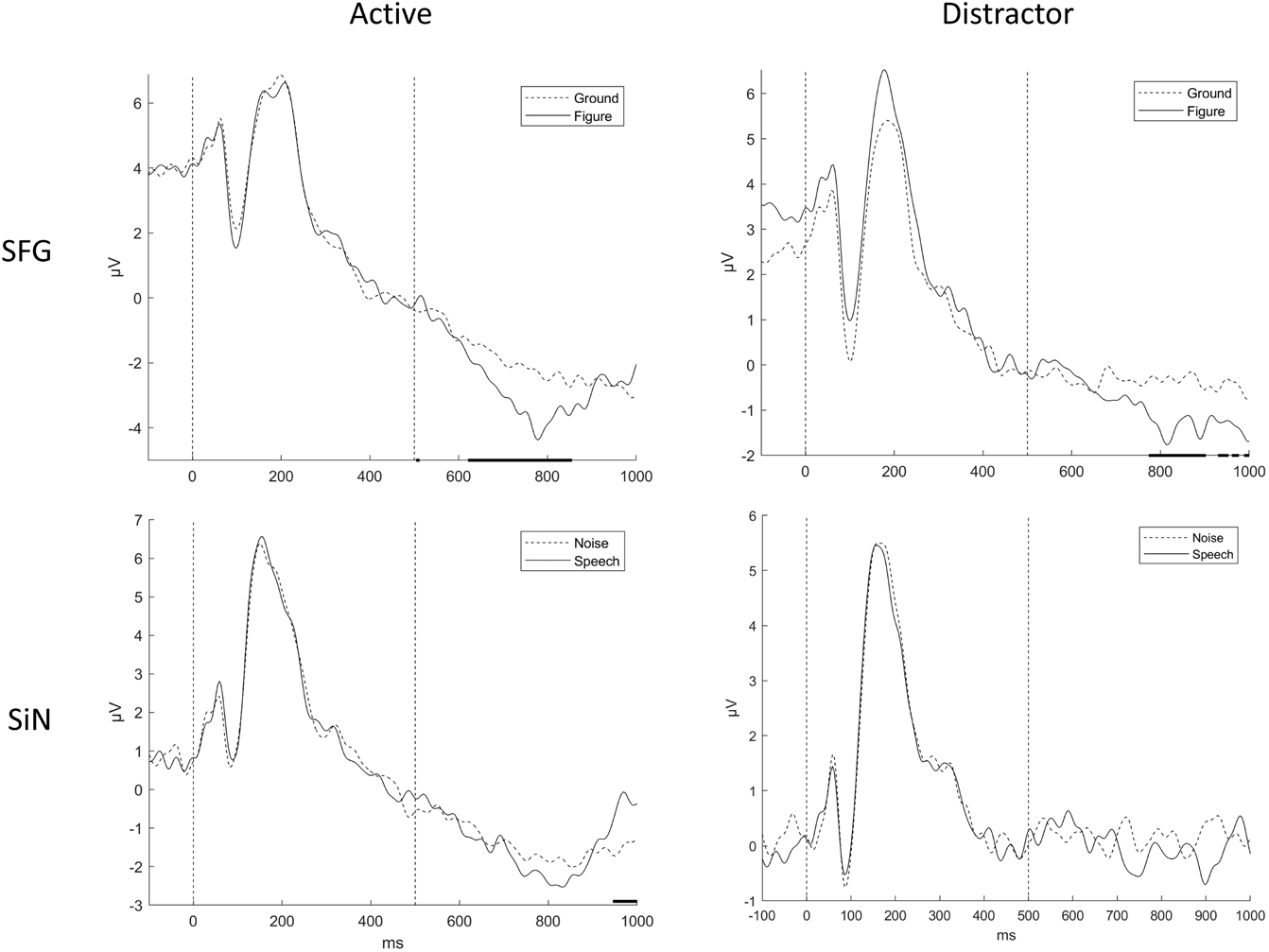
Group ERP waveforms at A1 on the active and distractor stochastic figure-ground test and the speech-in-noise test. Dotted lines signal auditory onset (0 ms) and stimulus onset (500 ms). Significance (p<0.05) based on non-parametric permutation cluster analysis is highlighted in black above the x axis.

### 2.3 Individual ERP Analysis

To evaluate the potential for clinical use, where group analysis is not possible, individual data were also examined (Figure 4), by taking the average difference between either figure and ground or speech and noise, over the time period 600 to 800 ms. On average, participants showed increased negativity when the target sound was present (figure or speech) (mean ± SD; active SFG: −1.09 ± 1.09; distractor SFG: −0.38 ± 1.09; active SiN: −0.27 ± 1.12; distractor SiN: −0.20 ± 0.10). This difference was robustly found across a majority of participants during the active SFG, as can be seen at the top of Figure 4, while SiN failed to elicit amplitude differences in over a third of participants. The separation of figure/ground and speech/noise is prominent for most participants. 15 out of 18 participants showed negative value for the amplitude differences of figure and ground in the active condition, 3 weakly showed the opposite pattern, and 3 participants showed very little effect of figure versus ground. The active condition showed a distinctive advantage over the distractor condition regarding the consistency of the activation pattern (10/18 had a negative figure-ground value), but separation is nevertheless evident for most participants (14/18) in the distractor condition. The SiN paradigm showed similar distribution, but around half of the individual data showed the opposite pattern compared to the group analysis in both conditions. The overall individual data and example waveforms from two selected participants are illustrated in Figure 4.

**Figure 4.**
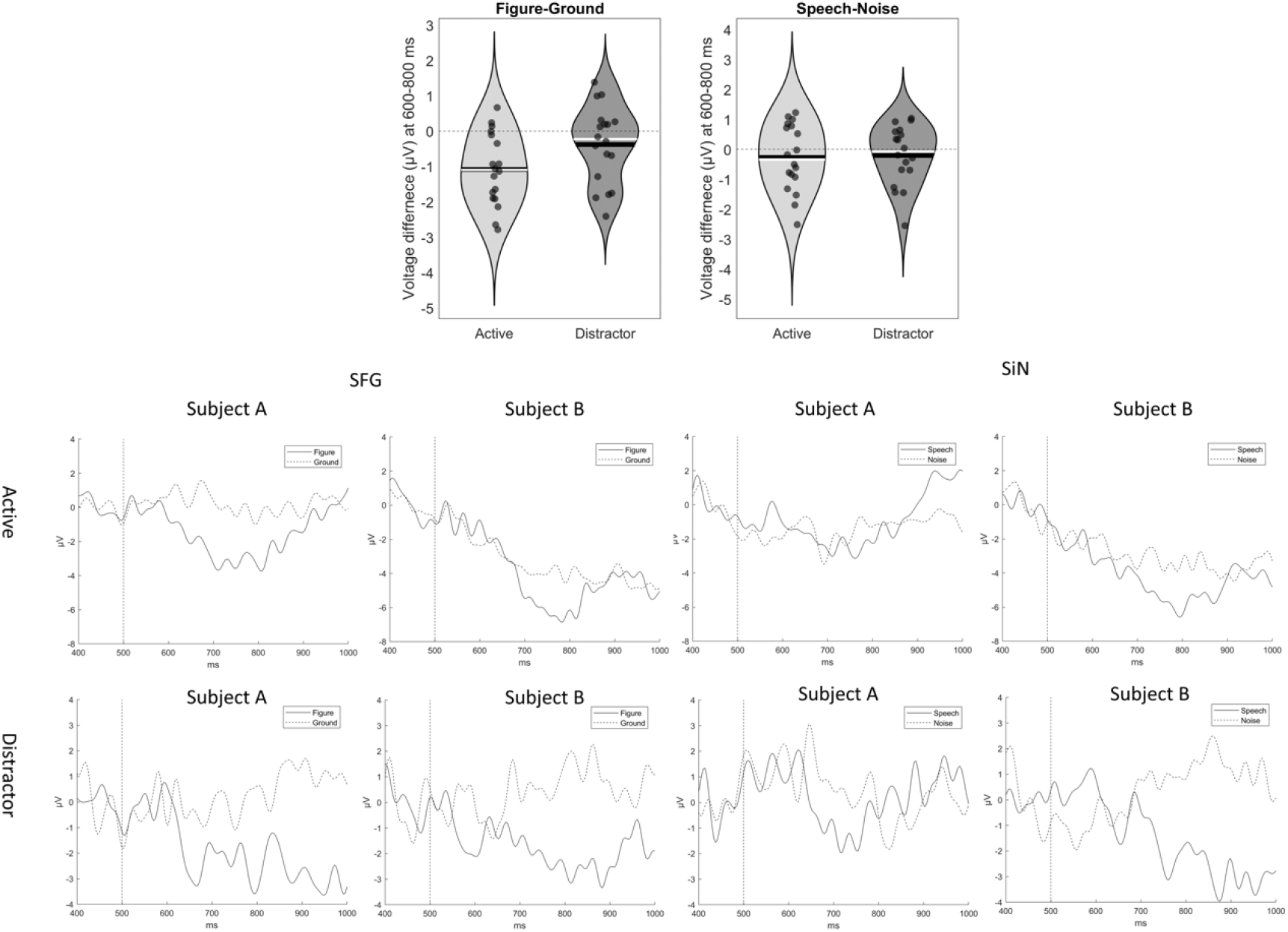
Individual data of all 18 participants. The top two violin plots show the distribution of the voltage differences of figure and ground over the time period 600 to 800 ms, and speech and noise in 18 participants. The mean and the median are highlighted in black and white, respectively. The bottom two rows are example waveforms of two typical participants.

The ANOVA test revealed a significant main effect of ‘Stimulus Type’ (F (1, 17) = 4.76, p=0.04, η_p_^2^= 0.22), which was due to a lower main amplitude difference for SFG than SiN (Table 2). The main effect of ‘Condition’ was also significant (F (1,17) = 9.25, p=0.007, η_p_^2^=0.35). The interaction between ‘Stimulus Type’ and ‘Condition’ was not significant (F (1,17) =1.23, p=0.28, η_p_^2^=0.07).

**Table 2.** Descriptive statistics of the EEG data. Descriptive statistics of the EEG data. They are speech minus noise and figure minus ground from left to right in active and distractor conditions (top-down).

|  | SiN (M/SD) | SFG(M/SD) |
| --- | --- | --- |
| Active | -0.27 (1.12) | -1.09 (1.01) |
| Distractor | -0.20 (0.10) | -0.38 (1.09) |

## 3. Discussion

The behavioural data demonstrated reliable task performance for all participants in both tasks, with a generally high d’ score. This shows that healthy-hearing people could easily detect the auditory target in these tests. When comparing the two active tasks, SFG did not show a significantly higher detection sensitivity (d’) than SiN, indicating a comparable SNR level. The visual d’s showed higher performance compared to the auditory tasks, which means that the visual distractor paradigm was robust in engaging participants’ attention. The audiogram did not show significant correlation with the outcome measures. This is likely due to the relatively small sample size and the small range of hearing ability from the normal hearing participants.

### 3.1 ERP Responses to Auditory Grouping

The hearing tests demonstrated robust EEG responses of figure and speech with a latency of around less than 200 ms in both active and distractor conditions. The figure evoked greater negativity over the vertex than when it was absent, which was also seen for the speech albeit with a weaker effect. The rapid figure-ground segregation, as well as the slow drift of the SFG responses, were also found in the MEG study (Teki et al., 2011), where the researchers observed shorter latencies for SFG compared to an EEG study by O’Sullivan et al. (2015). These responses are also consistent with the ORN reported by Tóth et al. (2016) in their EEG study. ORN is considered to reflect neural activity that occurs while actively segregating concurrent sounds (Alain et al., 2002). As the behavioural data have shown that the visual distractor in this experiment reliably engaged attentional resources, and the brain responses to SiN also exhibited a clear suppression of speech tracking under the distractor paradigm. Conversely, the persistence of figure detection responses under the SFG distractor condition indicates that spectrotemporal grouping could be a pre-attentive process. The SiN test also yielded a pattern of activation that was less consistent on individual analysis than for SFG. The SFG paradigm therefore could potentially provide a more robust neurophysiological measure for central grouping than the SiN test.

The topographic maps of SFG showed distinctive central negativity that is consistent with previous EEG work (Tóth et al., 2016) which localised the brain sources of the spectrotemporal grouping to the superior temporal gyrus and the inferior parietal sulcus, also in line with neuroimaging studies on SFG (Holmes et al., 2021; Teki et al., 2011). Furthermore, a cluster of central channels was revealed to be the major source of activation that powered the figure grouping, which supports the use of a single channel at the vertex for analysis. As the single channel analysis demonstrated very similar waveforms with minor differences in the statistically significant time points, and the recording setup as well as data analysis procedures are relatively simple, it is potentially a more optimal measure that could be adapted for clinical use.

The individual data showed that visible figure segregation could be seen in most participants and a majority of the participants showed a consistent activation pattern with the group-level ERP analysis. This means that the SFG paradigm could be used with EEG recording as a measure for auditory central grouping mechanism, and the results could be quantified by extracting a single metric (the average difference between 600-800 ms) from the EEG data and compared to 0. In contrast, the SiN paradigm in the current study did not exhibit reliable neural responses at either the group or individual levels. The ANOVA test showed that SFG also elicited significantly higher negativity compared to SiN suggesting that SFG is a more robust tool for neural responses to auditory grouping.

In conclusion, this study provides proof of principle for the utility of SFG as a complementary hearing test for SiN perception in clinical settings. It could reliably elicit individual behavioural and EEG responses that can easily be obtained in clinical settings with a single channel at the vertex. The visual distractor condition also showed group-level responses, indicating that SFG responses in EEG do not require any specific attention. Further studies are still required to produce a standardised clinical test, and additional steps still required also include studies in older populations, patients with hearing impairment, and performing correlations between SFG behavioural and EEG responses and clinical measures of speech in noise difficulty.

## Acknowledgements

We thank Phyllis Leung, Jerermy Pulmano, Violeta Ivanova, Holly Jenkins, and Ysanne deGraaf for helping us in the EEG data acquisition.

1 Abbreviation: stochastic figure-ground (SFG), speech-in-noise (SiN), pure tone audiogram (PTA), electroencephalography (EEG).

